# Targeted memory reactivation elicits temporally compressed reactivation linked to spindles

**DOI:** 10.1101/2024.09.12.612625

**Authors:** Mahmoud E. A. Abdellahi, Martyna Rakowska, Matthias S. Treder, Penelope A. Lewis

**Affiliations:** School of Psychology, Cardiff University Brain Research Imaging Centre (CUBRIC), Cardiff CF24 4HQ, United Kingdom; School of Engineering, Computer Science department, University College London (UCL), London NW1 2AE, United Kingdom; Faculty of computers and artificial intelligence, Cairo University, Giza 12613, Egypt; School of Computer Science and Informatics, Cardiff University, Cardiff CF24 3AA, United Kingdom

## Abstract

Memories reactivate during sleep, however the properties of such reactivation and its relationship to encoding strength and subsequent memory performance are not well understood. We set out to examine memory reactivations associated with a serial reaction time task (SRTT). 48 human participants performed the SRTT, and then slept in the lab while we deliberately induced reactivation in Slow Wave Sleep (SWS) using a Targeted Memory Reactivation (TMR) design. We detected reactivation after TMR cues using multiclass classification that adapted to sleep data by using sleep activity for training and wake activity for testing. We then examined the temporal properties of reactivation in relation to behavioural performance and sleep spindles. In keeping with the rodent literature, the observed reactivation was 3 to 20 times faster than waking activity. Furthermore, we report an inverted-U shaped relationship between TMR-related behavioural improvement and encoding strength, with very strong and very weak memories benefiting little from cueing while medium-strength memories benefit the most. Finally, reactivation was more frequently observed in trials with high sigma power, supporting the idea that sleep spindles are associated with memory reactivation during sleep. These findings bring us closer to understanding the characteristics of human memory reactivation after TMR, demonstrate when cueing is effective, and provide evidence for the positive relationship between the detectability of reactivation and memory consolidation.

## Introduction

We spend around one third of our lives asleep. During sleep, the brain is busy processing memories through replay or reactivation which is essential for memory consolidation (Diekelmann & Born, 2010; Rasch & Born, 2013; Ólafsdóttir et al., 2018; Wilson & McNaughton, 1994).

The active system consolidation (ASC) hypothesis (Diekelmann & Born, 2010) suggests that sleep is not merely a passive shelter for memories against interference. Instead, newly encoded memories repeatedly reactivate during slow wave sleep (SWS) and are thus transferred from the short-term store (hippocampus) to the long-term store (neocortex) in an ongoing process of memory consolidation. The ASC model (Rasch & Born, 2013) proposes a dialogue between neocortex and hippocampus in which slow oscillations (SOs) drive reactivation of hippocampal memories, with accompanying sharp wave ripples that are carrying reactivations nested into thalamo-cortical spindles. The model also suggests that spindles prime the cortex for reactivation-related plasticity by stimulating calcium influx into the dendrites of cortical pyramidal cells.

A technique called targeted memory reactivation (TMR) can be used to manipulate reactivation in sleep. In TMR, cues such as odours, sounds, or electrical shocks are associated with the learned material as a result of being presented during memory encoding or retrieval. Cues are then re-delivered during subsequent sleep and thereby thought to reactivate the cued memory (Hennevin & Hars, 1987). In humans, several studies have shown the benefits of TMR on memory consolidation for both declarative (Cairney et al., 2014; Fuentemilla et al., 2013; Rasch et al., 2007; Rudoy et al., 2009) and non-declarative memories (Antony et al., 2012; Monika Schönauer et al., 2014). Memory reactivation elicited via TMR can be detected using multivariate pattern classifiers and similarity analyses (Abdellahi et al., 2023a; Belal et al., 2018; Cairney et al., 2018; Schreiner et al., 2018; Wang et al., 2019). However, despite extensive research in the area, there are still a lot of gaps in our understanding of the characteristics of cued reactivation. Are such reactivations exact clones of wake activations, or do they differ in shape or duration? Does TMR favour certain memories based on their strength? How do sleep spindles relate to memory reactivation? And how does reactivation detection relate to consolidation? Here, we set out to answer these questions and thereby gain a better understanding of memory reactivation and TMR.

The reactivation-spindle connection is supported by Cairney and colleagues who showed that spindles mediate reactivation in human NREM sleep (Cairney et al., 2018). Additionally, a significant post-cue reactivation was observed in trials with high post-cue power in the spindle band (Wang et al., 2019), while enhancing spindles led to more consolidation (Lustenberger et al., 2016; Ngo et al., 2013). It has also been shown that hippocampal sharp-wave ripples are nested in the troughs of spindles (Staresina et al., 2015). Our current study will investigate how sleep spindles relate to reactivation.

In rats, memory replay during NREM sleep has been shown to have different temporal characteristics compared to wake, as it occurs from 10 to 20 times faster (Ji & Wilson, 2007; Lee & Wilson, 2002; Nádasdy et al., 1999). In wake, offline replay is thought to occur from 6 to 7 times faster than the actual task (Euston et al., 2007). Here, we set out to identify temporal compression during memory reactivation in human SWS using EEG data.

Sleep is thought to impact upon weakly and strongly encoded memories in very different manners. On the one hand, sleep provided a preferential benefit to weaker memories during encoding in several studies (Cairney et al., 2016; Creery et al., 2015; Drosopoulos et al., 2007; Lo et al., 2014), and motor skill learning itself showed greater gains across sleep for objects with lower pre-sleep performance (Kuriyama et al., 2004). Furthermore, consolidation of weakly encoded items was associated with fast sleep spindles density (12.5-16 Hz) and spindles that were coupled with SOs predicted weak memory consolidation (Denis et al., 2021). On the other hand, participants who encoded memories most strongly have also been shown to benefit from sleep the most (Tucker & Fishbein, 2008). On the back of this apparently conflicting literature, Stickgold proposed that consolidation of memories follows an inverted U-shaped distribution, such that memories with very high and very weak strength benefit little from sleep, while memories with average strength benefit the most (Stickgold, 2009). Given this literature, we set out to more closely examine how encoding strength influences consolidation using TMR.

We used a serial reaction time task (SRTT), which is known to be sleep sensitive (Born & Wilhelm, 2012; Spencer et al., 2006) and also sensitive to TMR in non-REM sleep (Cousins et al., 2014, 2016), (see figure 1 for experimental design and supplementary figure 1) to investigate the characteristics of cued reactivation. In the SRTT, participants saw an image on one of four quadrants of the screen and simultaneously heard a distinct sound that was associated with that image during encoding. We then distinguished between reactivation of four distinct memories after TMR cues by directly relating wake and sleep EEG in 48 participants. We introduce a classification pipeline in SWS that uses sleep activity for the training of classifiers and wake activity for testing, and which allows classifiers to adapt to sleep features that are related to reactivation when adjusting their weights. We then studied the differences between the activity of wake and sleep via machine learning models. We found that while the detected reactivation pattern mimics wake activation, it is temporally compressed 3 to 20 times compared to wakeful task activation. We also show that TMR selectively benefits memories that are not very strongly or weakly encoded. Finally, our results support prior work by showing that high post-cue sigma power is a marker for reactivation.

**Figure 1:**
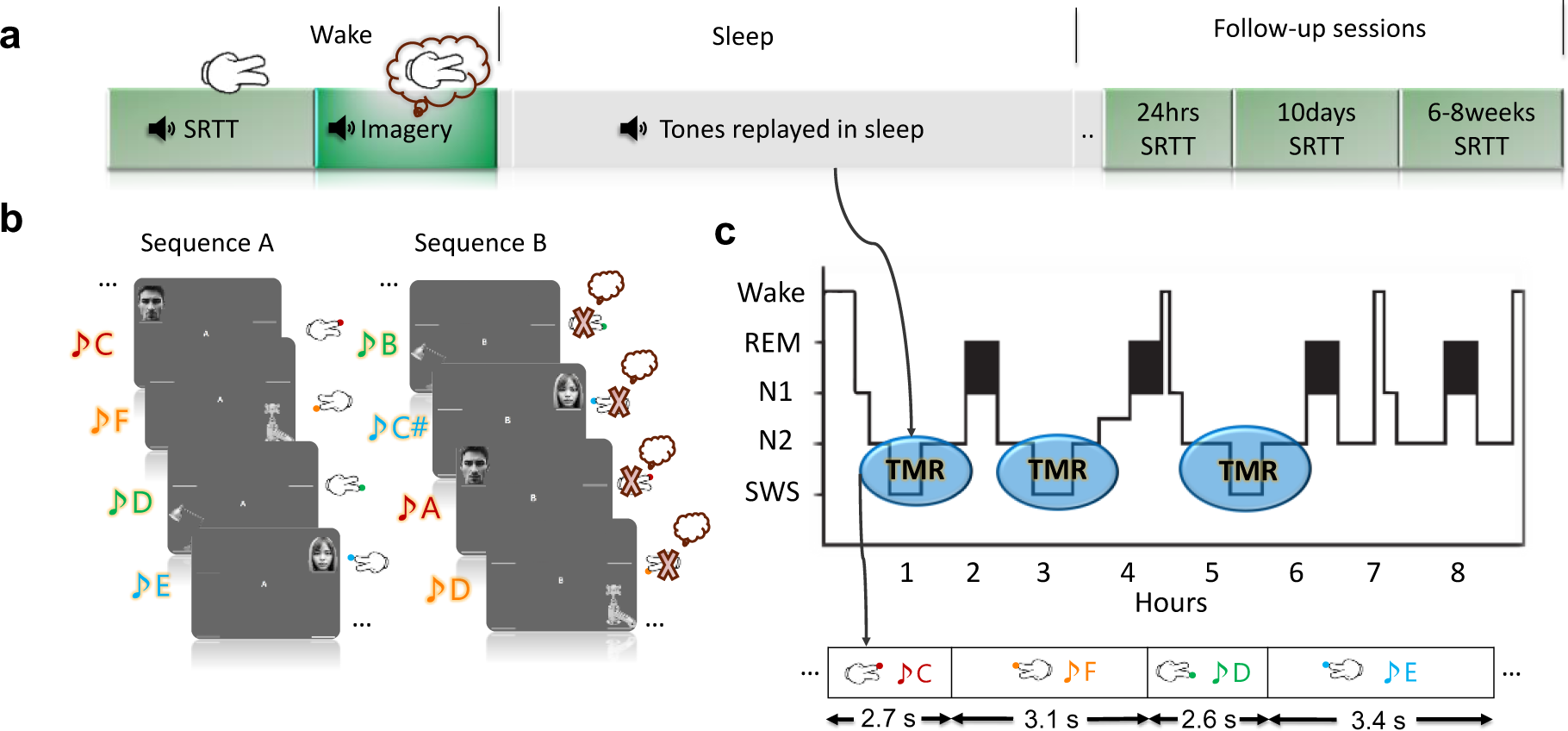
Study design. **a)** We analysed sleep and wake data from 48 participants. Participants first performed a serial reaction time task (SRTT), followed by a motor imagery task, both with the EEG headcaps on. Subsequently, they went to sleep and TMR was carried out in NREM sleep, as shown in panel c. After that, the participants were tested on the SRTT in three follow up sessions. **b)** In the SRTT, four images are presented in two different sequences. Each image is accompanied by a specific tone (different for each sequence) and requires a specific button to be pressed. In the imagery task, participants view the same sequences of images but only imagine they are pressing the buttons without any actual movements. This motor imagery task served as a clean template for characterising wake pattern and was later used in classification. **c)** TMR took place in NREM sleep with jittered intertrial intervals between 2500ms and 3500ms. Each sequence was followed by a 20-second pause.

## Results

### TMR benefits cued sequence

Studies on the SRTT have shown a positive effect of TMR on consolidation (Cousins et al., 2014, 2016; Koopman et al., 2020). Here, we tested TMR-dependent consolidation by comparing SRTT performance in the follow-up sessions between cued and un-cued sequences (see the methods section for details). We found a benefit for the cued sequence as compared to the un-cued sequence across follow-up sessions (Wilcoxon signed rank test, n = 41, p = 0.016, z = 2.42, figure 2a). This shows the positive effect that TMR has on memory improvement.

**Figure 2:**
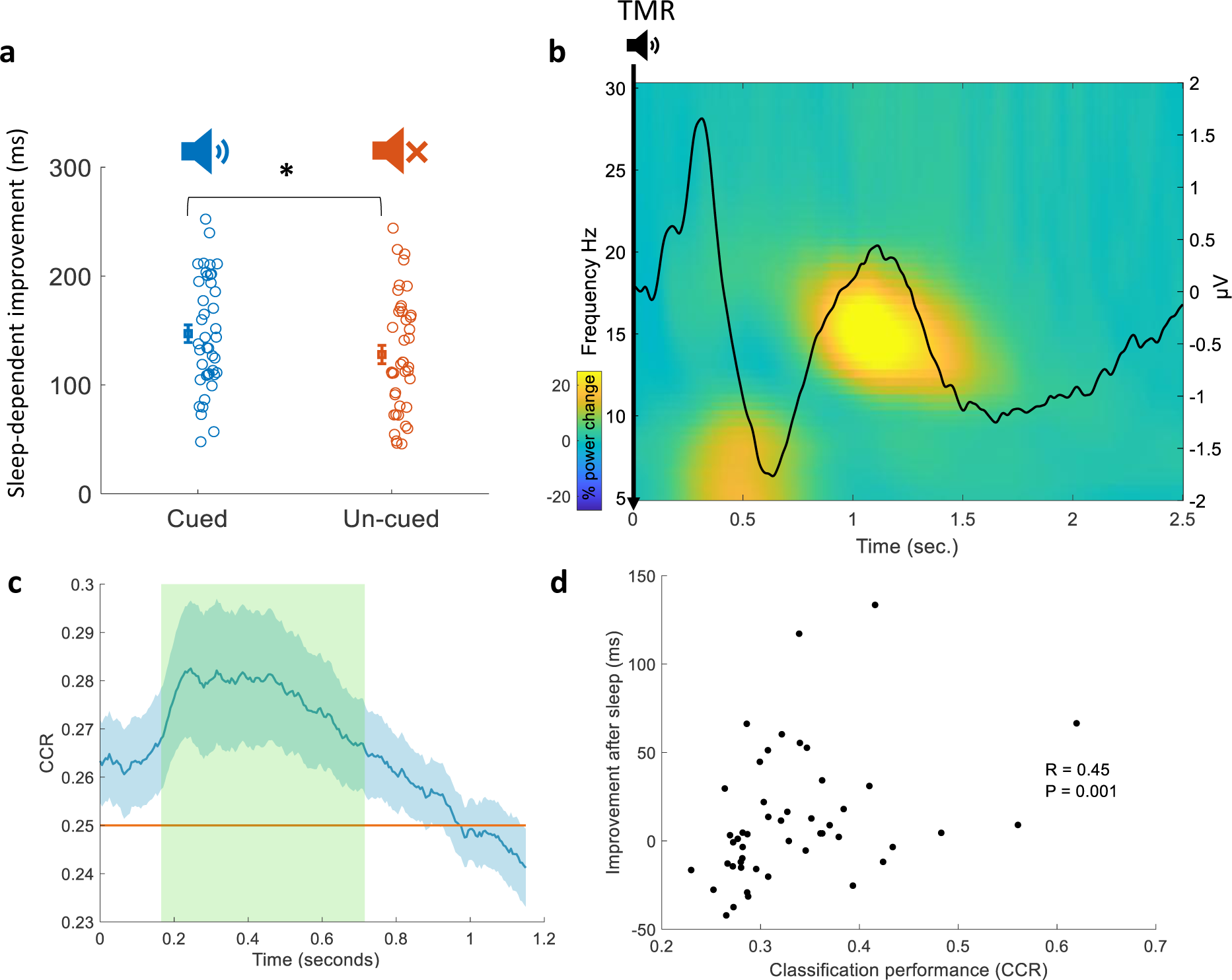
**a)** Behavioural improvement is significantly higher for the cued sequence compared to the un-cued one (Wilcoxon signed rank test, n = 41, p = 0.016, z = 2.42) indicating that TMR benefited the cued sequence. **b)** Time-frequency and ERP analyses using sleep data from all participants (n = 48). Power percentage changes from the baseline period [-0.3 -0.1] sec. are shown with colours. The solid black line represents the average results of all ERP analyses from all participants (n = 48). **c)** TMR elicited detectable reactivation. A linear classification shows a significant correct classification rate (CCR) compared to chance level of 0.25, this effect is explained by a cluster (green shaded area, n = 48, p = 0.026) after correcting using cluster-based permutation. **d)** Classification performance (CCR) correlated positively with memory improvement immediately after sleep (Spearman r = 0.45, p = 0.001, n = 48), a partial correlation controlling for pre-sleep behavioural performance also showed a significant correlation (Spearman r = 0.38, p = 0.009, n = 48).

### Elicited response after TMR cues

TMR has been shown to elicit a distinguishable oscillatory pattern that is apparent in the time-frequency representation as well as ERP analysis. We looked at the TMR-elicited response in both time-frequency and ERP analyses using a similar approach to (Cairney et al., 2018). As presented in figure 2b, EEG response showed an increase in theta band followed by an increase in sigma band, with the latter starting about one second after TMR onset. Furthermore, ERP analysis showed a small increase in ERP amplitude immediately after TMR onset, followed by a decrease in amplitude 500ms after the cue. These findings demonstrate that TMR was effectively eliciting a response, thus confirming that our TMR cues were being processed by the brain.

### Memory encoding activity during wake re-emerges in sleep after TMR cues

Several different methods for detecting memory reactivation have been adopted in the literature, some of which discriminated cued categories within sleep without the inclusion of wake (Cairney et al., 2018; M. Schönauer et al., 2017), while others selected features that showed high discrimination in wake (Wang et al., 2019). Our previous method directly relates wake and sleep activity using machine learning classifiers, but those classifiers were trained on wake and tested on sleep (Abdellahi et al., 2023a). We have now improved our method so that the classifiers pay attention to features present in sleep that are related to reactivation. We did this by building a machine learning model that was trained with the sleep data occurring after each TMR cue and tested during wakeful imagination of the trained task. This pipeline allows classifiers to weigh the features according to those present in sleep rather than weighing features according to those present in wake which could be dominated by effects that are absent from sleep. This also allows our linear classifiers to see the noise of sleep data represented in the within-class covariance.

Furthermore, unlike our prior work (Abdellahi et al., 2023a), where we classified only whether reactivation relates to imagined use of the right and left hand, in this current project we trained linear discriminant analysis (LDA) classifiers on our multi-class SRTT with each finger representing a class (4 classes in total, 2 fingers per hand). We used sleep data from 58 EEG channels and principal component analysis (PCA) to unravel the dimensions/principal components with highest variance. PCA can be used to reduce dimensionality and reduce overfitting and has been adopted in several studies (Griffiths et al., 2021; Higgins et al., 2021; Peyrache et al., 2010; Schreiner et al., 2021; Tingley & Peyrache, 2020). We then transformed the data using the principal components that explained the highest variance (see methods). The same transformation was done on wake data such that sleep and wake were transformed and the channels dimension became principal components.

Given that the task involved a sequence of trials in a fixed order, we were concerned that the brain might prepare responses in advance of the TMR cue. We therefore jittered the intertrial intervals between the TMR cues to eliminate this possibility. Trials therefore varied in durations by a maximum variation of one second between the shortest trial (2500ms) and the longest trial (3500ms). Given the uncertainty of the timing of reactivation, and the fact that it could sometimes happen after 2500ms., we included all of the temporal information of the sleep data into our classification model by using time points as observations (see methods). Sleep data was then used to train a LDA classifier, and this classifier was applied to wake at every time point after the sound cues, giving a classification performance (correct classification rate, CCR) at every time point in wake. Classification performance was significantly above chance level (figure 2c, significant effect is explained by the cluster with the green shaded area, p = 0.026), this shows that memory reactivation can be identified by our classification models.

### The strength of reactivation predicts memory consolidation

We wanted to test whether the elicited reactivation in sleep predicts the extent of TMR-dependent benefit right after sleep (24 hours). To this end, we conducted a Spearman correlation between the classification performance (CCR) and cueing benefit right after sleep (reaction time for non-reactivated sequence – reaction time for reactivated sequence). This showed as strong positive relationship (Spearman r = 0.45, p = 0.001, n = 48), figure 2d, supporting the idea that the reactivations detected by our classifiers underpin cueing benefit. To control for the effects of pre-sleep performance during encoding, we also conducted a partial correlation between classification performance and improvement right after sleep (Spearman r = 0.38, p = 0.009, n = 48), see methods. This revealed that the strength of reactivation positively predicts consolidation, again supporting a functional role for our detected reactivation.

### Memory reactivation in SWS is temporally compressed compared to wake

We next tested whether sleep reactivation mimics the shape and duration of wake activation by performing an analysis of compression and dilation. In this analysis, we fixed the length of wake trials and progressively changed the length of sleep trials. We used a ratio (length of sleep trial / length of wake trial) to indicate the temporal ratio between sleep and wake duration. Thus, a ratio of less than one indicates compression, a ratio of exactly one indicates no compression or dilation, and a ratio of greater than one indicates dilation. For every ratio, we applied a sliding window approach where we took sleep windows according to the ratio and then resized them to match the length of wake trials. Afterwards, we trained a classifier on sleep and tested it on wake to see if the sleep reactivation pattern was similar to wake at the given ratio (see methods). Our results indicate that sleep reactivation is compressed compared to wake, and this compression is 3 to 20 times faster than in wake. This result is in-line with the rodent literature, suggesting faster replay compared to wake activity (Euston et al., 2007; Ji & Wilson, 2007; Lee & Wilson, 2002; Nádasdy et al., 1999).

### Does TMR favour weakly encoded memories?

We were also interested to determine if the benefits of TMR depend upon the strength of the encoded memory. To do this, we examined pre-sleep reaction times, taking the best performance blocks as our measure of performance (see methods), and used them to rank participants from fastest to slowest. We then looked at the sleep-dependent improvement for different participants, that is, the difference in performance in the follow-up sessions between cued and un-cued sequences. This analysis revealed that neither strongly encoded (very fast) nor weakly encoded (slow) memories benefit from TMR. Instead, the biggest TMR-related improvement appeared in participants with average reaction times (figure 4a). Significant points in figure 4a also survived a cluster-based permutation test at p<0.0001 (see methods for details). This shows that the relationship between encoding and TMR benefit follows an inverted U-shape as shown in figure 4a, which is in-line with the literature supporting such a relationship (Stickgold, 2009).

**Figure 3:**
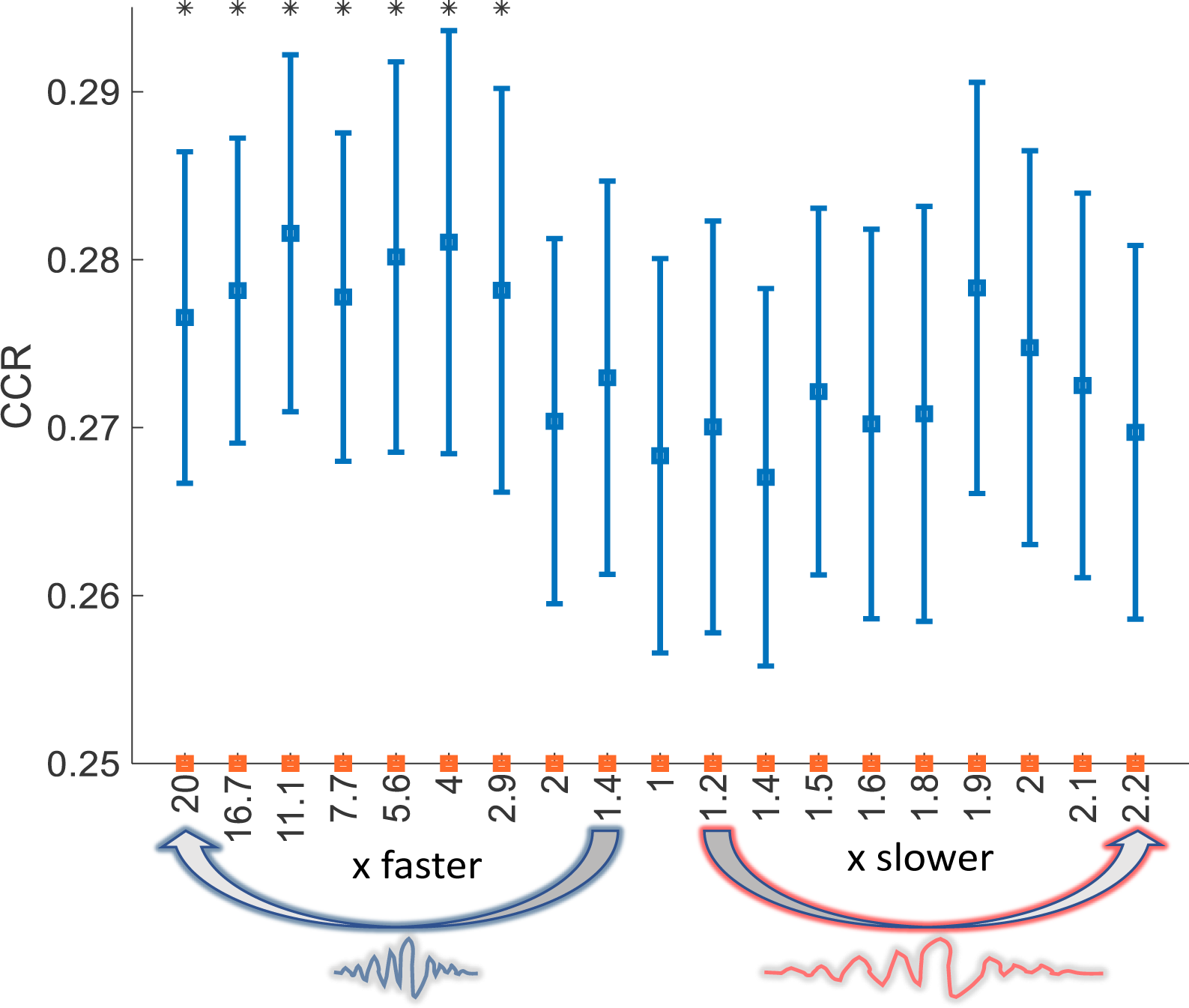
Analysis of temporal compression shows that reactivation is faster than wake pattern. The x-axis represents how much (x) faster or slower sleep reactivation was compared to wake, the y-axis represents correct classification rate (CCR). Significant results (p < 0.05) are marked by asterisks.

**Figure 4:**
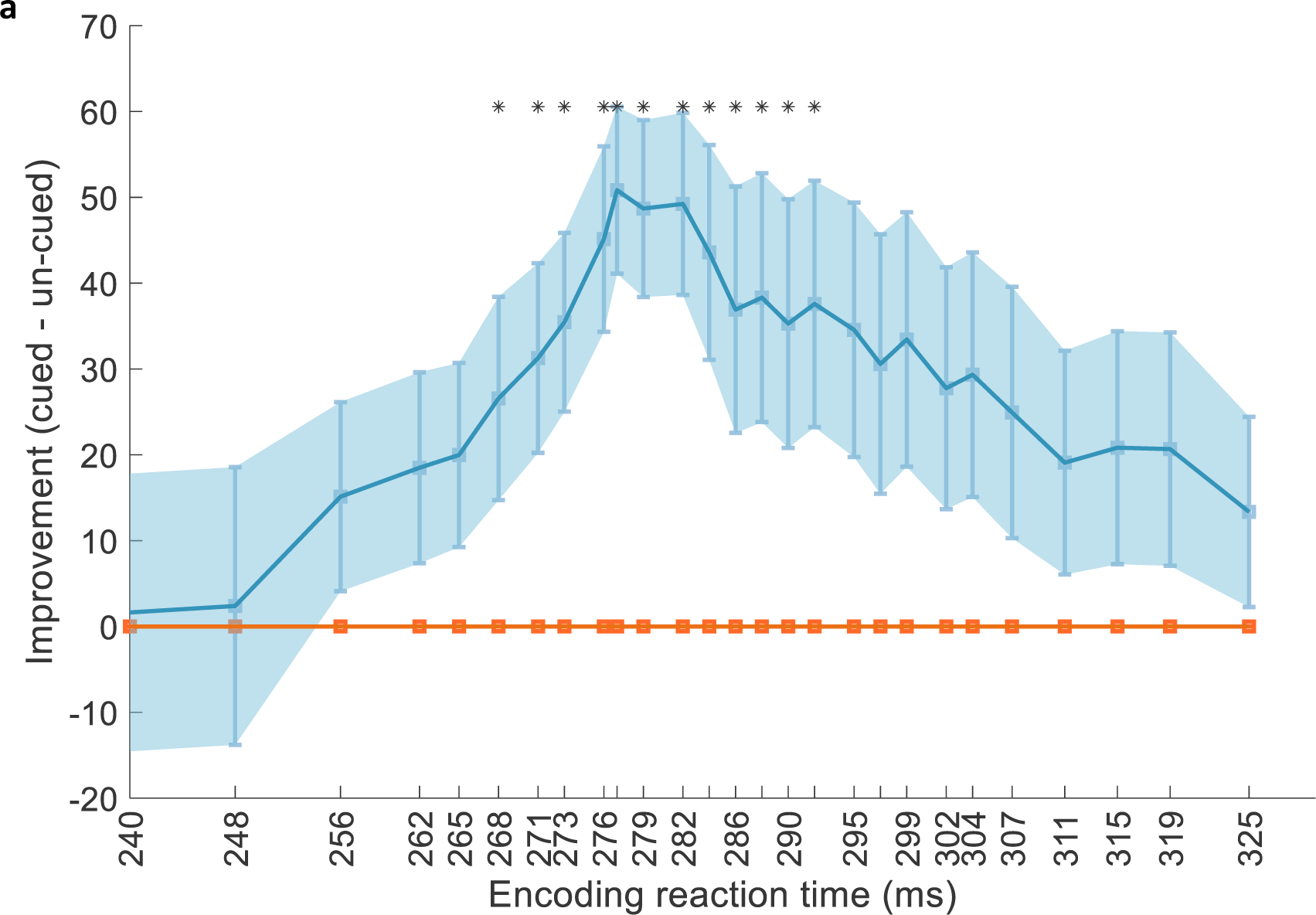
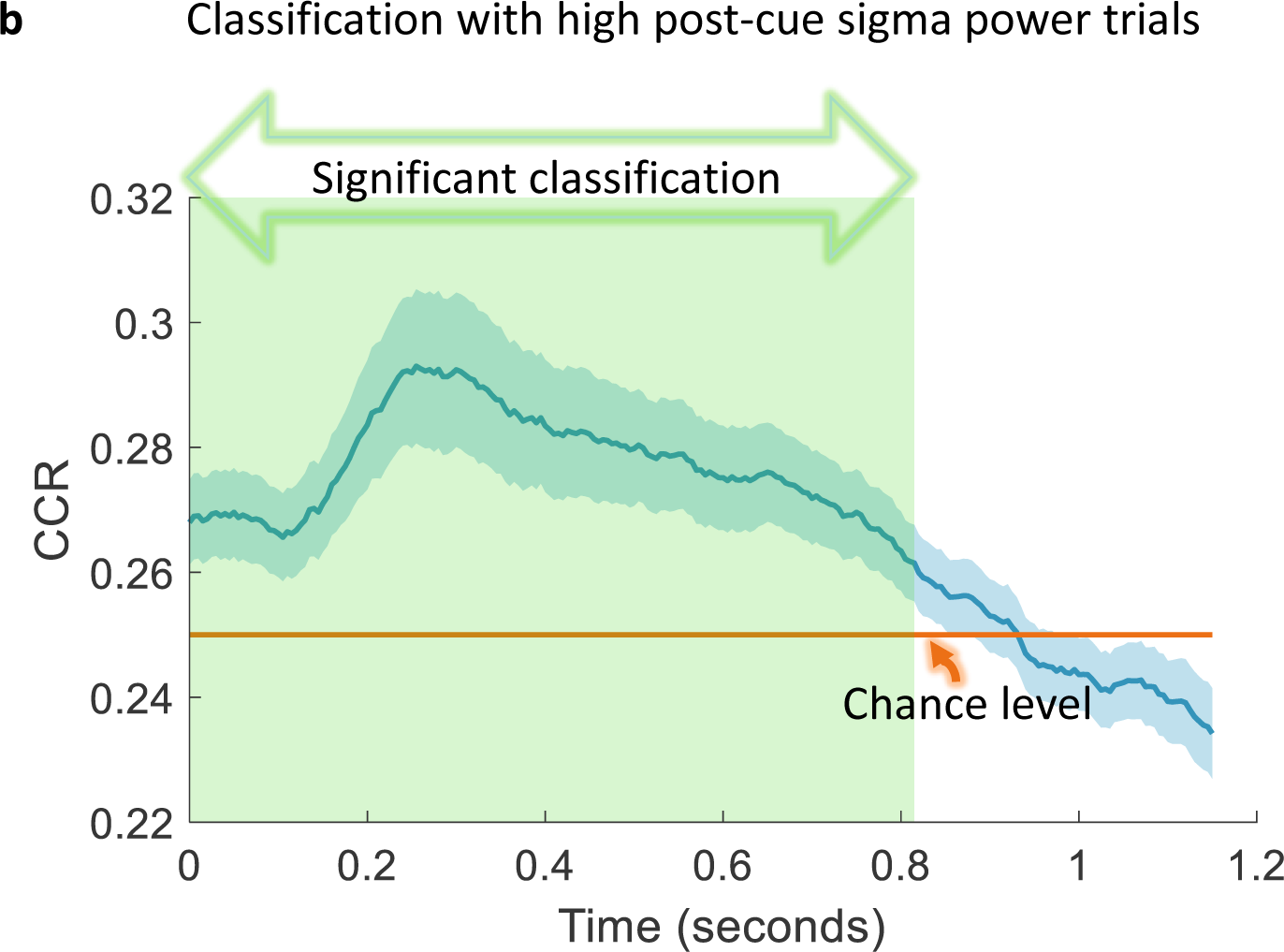
**a)** The relationship between encoding strength and sleep-dependent consolidation follows an inverted U-shape where very strongly/weakly encoded memories benefit very little from TMR. Significant points (p < 0.05) are indicated by asterisks and middle points that are significant also survived cluster-based permutation and showed a significant effect explained by a cluster (p < 0.0001). **b)** Classification using sleep trials with high post-cue sigma power [11 16]Hz shows significant classification performance explained by a cluster (green shaded area, p = 0.001). CCR, correct classification rate.

### Spindles hallmark reactivation

Wang and colleagues, who examined TMR cued NREM reactivation during a similar task showed that trials with high post-cue sigma power [11 16] Hz, were more likely to involve detectable reactivation (Wang et al., 2019). Here, we report the same relationship. After performing a median split on sigma power, similar to their analysis, we found that only trials with high post-cue sigma power showed evidence of reactivation (significant effect explained by a cluster p = 0.001, figure 4b) compared to chance level. This supports the idea that high post-cue sigma power acts as a marker for reactivation. Interestingly, classification of these high-sigma trials was also significant when compared to classification using low sigma power trials (significant effect explained by a cluster p = 0.022, supplementary figure 2).

## Discussion

We examined the temporal characteristics of the reactivation of individual finger representations associated with a SRTT and provide direct evidence that reactivation happens faster than the original experience during wake. We also show that memories primarily benefit from TMR when they are encoded neither very strongly nor very weakly, but with a medium strength. Finally, our results support earlier work suggesting that sleep spindles provide a marker of reactivation.

Some studies used only sleep data in their classification pipelines to show evidence for the reprocessing of memories during sleep (Cairney et al., 2018; M. Schönauer et al., 2017). Others performed within sleep classification with features selected from wake data (Wang et al., 2019) or by relating wake to optimal sleep lags (Belal et al., 2018). Here, we directly related neural responses in sleep to those during the imagery task in wake by training classification models on sleep observations and applying them on wake. This direct sleep-wake relationship means that our models will not mistakenly classify sleep EEG noise as reactivations. Thus, our linear classifiers can adapt to sleep and adjust their feature weights according to sleep patterns. This also enables our models to see sleep noise represented by within-class covariance matrices and adapt to it. We successfully used this approach in classifying memory reactivation after TMR in human REM sleep (Abdellahi et al., 2023b). To further elucidate the wake-sleep relationship, we used jittered inter-trial delays, thus preventing periodic oscillations from affecting the training of our models. Given that the finger-tapping task is a sequence, if we were to use fixed inter-trial delays the brain could have predicted and reactivated the contents of the upcoming TMR before it has actually been presented. Our jittered cues avoided this possible predictability.

Several rodent studies have tackled the question of temporal compression of reactivation. Findings show that cell firing happens at a faster rate during replay compared to the original experience (Davidson et al., 2009; Euston et al., 2007; Ji & Wilson, 2007; Lee & Wilson, 2002; Nádasdy et al., 1999). Collectively, replay has been observed at different rates, ranging from 6 to 20 times faster than the waking experience. Given that the mentioned studies show results from firing activity in rodents and here we use EEG signals from human participants who encoded multiple memories, we used sleep-wake classifiers to address the question of compression. Our results are in-line with the literature, suggesting that reactivation happens at a rate that is around 3 to 20 times faster than wake.

It has been proposed that memories are transferred into a long-term store via repetitive reactivation (Diekelmann & Born, 2010). According to this view, there is a dialogue between the hippocampus and the neocortex wherein cortical SOs drive thalamo-cortical spindles. Ripples and their associated reactivations are nested in the troughs of these spindles, which emphasises the importance of sleep spindles and ripples in the reprocessing of memories. Several papers have shown a direct relationship between memory reactivation and spindles in which spindles marked reactivation (Cairney et al., 2018; Wang et al., 2019). Moreover, Zhang and colleagues provided direct evidence that human memory replay happens during ripple events using intracranial EEG and similarity analysis (Zhang et al., 2018). We provide direct evidence of reactivation being marked by spindles, thus supporting the hypothesis that reactivation occurs during ripple events. This could explain why it is compressed in time. Indeed, the compression of 3 to 20 times observed in our data means that reactivations happen for a duration of 57ms to 383ms which could support the speculation that ripples can carry reactivations, since they are characterised by 50 to 100ms of high frequency activity (Ylinen et al., 1995). Despite the technical limitations of directly estimating ripple events in human cortical EEG, our temporal compression analysis helps to unravel the footprint of ripples and the impact they have on the temporal characteristics of the detected reactivation. Along with spindle analysis, this evidence fits well with the idea of spindle-ripple events as a hallmark for reactivation.

The strength of the encoded memories during wake could be a factor in determining the mechanism by which memories are prioritised for consolidation. Intuitively one may think that very strong memories do not have much room for improvement and thus cannot benefit from sleep-dependent consolidation. Indeed, several studies have now shown that weakly encoded memories benefit from sleep-dependent consolidation (Cairney et al., 2016; Creery et al., 2015; Drosopoulos et al., 2007; Lo et al., 2014). However, at least one study has shown that sleep-dependent consolidation favours strongly encoded memories (Kuriyama et al., 2004). Furthermore, it has also been suggested that consolidation may follow an inverted U-shape, such that very weak and very strong memories do not benefit from sleep (Stickgold, 2009). Our data support this latter suggestion by showing a TMR benefit for medium strength memories, and very little TMR benefit for very weak and very strong memories (figure 4b). There is evidence that cortical networks are being “tagged” during encoding and that this tagging is needed for memory consolidation (Lesburguères et al., 2011). Lewis and Bendor proposed a mechanism for how TMR could bias replay (Lewis & Bendor, 2019), proposing that during behaviour, cortical networks related to a memory are being tagged and their priority for consolidation is determined in a cortico–hippocampal–cortical loop. Here, we suggest that behaviour during encoding somehow tags memories as a priority for consolidation if they were encoded neither very strongly nor very weakly.

Together, our findings show that slow wave sleep reactivation in humans occurs faster than activation during the task. We also show that TMR does not favour very weakly or very strongly encoded memories for consolidation, but those of a medium strength instead. Finally, we support prior work showing that spindles are hallmarks for reactivation and that the detected reactivations predict memory consolidation which reflects their functional significance. Overall, we describe new characteristics of reactivations and how they relate to wake. We also introduce a new method for detecting SWS reactivation by training classification models with sleep EEG and testing them on wake data.

## Methods

### Participants

We collected EEG and behavioural data from human participants (n = 48) (25 females, age mean ±SD: 19.9 ±1.4; 23 males, age: 20.8 ±2.1). The number of participants was reduced from 56 as some of them were excluded for technical problems during recording of sleep. Participants completed the SRTT before sleep and during three follow up sessions, the first one was after the night of stimulation (24 hours), the second after 10 days later, and eventually the final session after 6 to 8 weeks. All participants were right-handed with no prior knowledge of the SRTT. All participants had normal or corrected-to-normal vision, normal hearing, and no history of physical, psychological, neurological, or sleep disorders. Responses in a pre-screening questionnaire reported no stressful events and no travel before commencing the study. Participants did not consume alcohol or caffeine in the 24 hours prior to the study or perform any extreme physical exercise or nap. This study was approved by the School of Psychology, Cardiff University Research Ethics Committee, and all participants gave written informed consents.

### Experimental design

Participants completed the SRTT adapted from (Cousins et al., 2014). Participants learned two 12-item sequences, A and B (A: 1 2 1 4 2 3 4 1 3 2 4 3 and B: 2 4 3 2 3 1 4 2 3 1 4 1). Sequences had been matched for learning difficulty; both contained each item three times. Sequences were presented in blocks and each block contained three repetitions of a sequence. The blocks were interleaved so that a block of the same sequence was presented no more than twice in a row. There were 24 blocks of each sequence (48 blocks in total), and each block was followed by a pause of 15 seconds during which feedback on reaction time (RT) and error-rate were presented. After the 48 blocks of sequences A and B, participants performed four blocks of random sequences. They contained the same visual stimuli, two of these blocks were paired with the tone group of one sequence (reactivated in sleep), and the other two with the tone group of the other sequence (not reactivated). Participants were aware that there were two twelve-item sequences, and each sequence was indicated with ‘A’ or ‘B’ appearing centrally on the screen, but participants were not asked to learn the sequences explicitly. Counterbalancing across participants determined whether sequence A or B was the first block, and which of the sequences was reactivated during sleep. Each sequence was paired with a group of pure musical tones, either low tones within the 4th octave (C/D/E/F) or high tones within the 5th octave (A/B/C#/D). These tone groups were counterbalanced across sequences. For each trial, a 200ms tone was played, and at the same time a visual cue appeared in one of the corners of the screen. The location indicated which key on the keyboard needed to be pressed as quickly and accurately as possible: 1 – top left corner = left shift; 2 – bottom left corner = left Ctrl; 3 – top right corner = up arrow; 4 – bottom right corner = down arrow. Participants were instructed to keep individual fingers of their left and right hand on the left and right response keys, respectively. Visual cues were neutral objects or faces, used in previous studies (Cousins et al., 2014), which appeared in the same position for each sequence (1 = male face, 2 = lamp, 3 = female face, 4 = water tap). After responding to the visual cues with the correct key press an 880ms inter-trial interval followed.

After completion of the SRTT, participants were asked to do the same task again, but were instructed to only imagine pressing the buttons. Motor imagery (IMG) consisted of 30 interleaved blocks (15 of each sequence), presented in the same order as during the SRTT. Each trial consisted of a 200ms tone and a visual stimulus which was presented for an 880ms followed by a 270ms inter-trial interval. There were no random blocks during the imagery task and no performance feedback was presented during the pause between blocks. We collected the SRTT data during three sessions after the stimulation night, with one the next day (24 hours) after performing the task and spending the night in the lab, the second one after 10 days and the third after 6 to 8 weeks. During the night of stimulation cues were presented in during NREM sleep with the continuous supervision of experiments and data scored as N3 was the one included in the analysis. Inter-trial intervals were jittered between 2500ms and 3500ms, as demonstrated in figure 1. Stimulation was paused with any signs of arousals until the experimenters observe approximately three 30-second epochs with stable NREM sleep. In the follow up sessions (24 hours, 10 days, and 6 to 8 weeks) after the task, participants were asked to perform the SRTT again. Eventually, in the last session, they were asked if they remember the locations of images of the two sequences in order to see if one sequence is recalled better than the other one. Motor imagery data set of each participant was used to classify the brain activity without movement artifacts.

### Behavioural improvement

We measured the behavioural improvement after sleep in three different sessions, the first was after sleep and the second after 10 days and the third after 6-8 weeks. Some participants were excluded from the analysis because they dropped out and did not come to the follow ups, thus, the number of participants in this analysis was 41 participants. We were interested in the aggregated effect of TMR across these sessions. For every session, all 24 blocks containing the reaction times for a sequence were aggregated and the blocks with the best performance among them were kept based on the 95 percentiles of performance values. Thus, the fastest 5 percentiles of data are used from every session and the median of post-sleep sessions was calculated. The same procedure was conducted for pre-sleep session where the fastest 5 percentiles of blocks were used as the pre-sleep performance measure. Afterwards, we determined the improvement as (pre-sleep – post-sleep), thus a high value would reflect big improvement. We then tested for the difference between the improvement for the reactivated and the non-reactivated sequence using a Wilcoxon signed-rank test.

### The relationship between encoding behaviour and consolidation

We performed an analysis on encoding performance and subsequent improvement to study how they relate and whether the prioritisation of certain memories comes down to the encoding salience. In this analysis, we sorted participants based on their performance during encoding. Thus, participants with the lowest reaction times represent the fastest performers. We then took a cluster that included 15 participants based on the reaction times as we are interested in the general relationship between encoding and improvement rather than effects from individual participants. Also, with this approach we were able to evaluate the improvement and whether it is significant for each cluster. We thus included 15 participants in each iteration based on the sorted reaction times, such that the first point in figure 4a includes the 15 participants with the lowest reaction times (fastest). We then shifted this cluster by one and thus included a participant with a higher reaction time before sleep and removed the participant with the lowest reaction time. We proceeded until the participant with the highest reaction time. The difference between the improvement for the cued and the un-cued sequence for each point was then plotted and tested with a Wilcoxon signed-rank test. We considered that grouping the participants will make the data points dependent instead of being independent, so we repeated the statistical tests but using cluster-based permutation and found significant effect explained by a cluster which contained the significant points from the Wilcoxon signed-rank test.

### The relationship between reactivation strength and memory consolidation

We performed a correlation analysis between the classification performance of reactivation and memory improvement after sleep. Memory improvement for each participant was measured as the difference between the reaction time of the un-cued and the cued sequence, which reflects the cueing benefit. To measure the relationship between reactivation and the direct cueing benefit we used the follow up session that came after sleep. The strength of memory reactivation was determined by the maximum classification CCR value for each participant. In this partial correlation, we controlled for the effects of the encoding session reaction times. Blocks of behavioural reaction times were aggregated into one value for each participant in the same way we calculated the behavioural improvement by keeping the fastest 5 percentiles of performance values and then taking their median.

### EEG recording

The current study uses EEG from human participants. EEG was collected using 64 actiCap active electrodes with 62 channels on the scalp including the reference electrode at CPz and ground electrode at AFz. Two electrodes were used on the left and right sides above and below the eyes for collecting electrooculography (EOG) signals and two electrodes on the right and left sides of chin for collecting the electromyography (EMG). Data were collected at 500Hz and 250Hz and subsequently resampled to 200Hz for all EEG analysis. Sound cues were delivered during NREM sleep.

### EEG cleaning

EEG cleaning consisted of filtering and outliers’ rejection based on statistical measures. EEG data were band-pass filtered (0.1 to 30Hz) and centred. For sleep data, sleep was scored manually and only the trials in the epochs scored as N3 were used in this work. Afterwards, we removed trials representing outliers based on statistical measures (variance, max, min) extracted for every trial and every channel. A trial is compared to all trials and considered as an outlier if it was higher than the third quartile + (the interquartile range *1.5) or less than the first quartile - (the interquartile range*1.5) in more than 25% of channels. If a trial was bad for <25% of channels it was interpolated using neighbouring channels with triangulation method in Fieldtrip. Furthermore, because the task is motor-related we defined a number of channels around the motor area (C6, C4, C2, C1, C3, C5, CP5, CP3, CP1, CP2, CP4, and CP6) and a trial was rejected if it is bad on >25% of these channels otherwise bad channels are interpolated and the trial was kept.

### Detecting memory reactivation with multivariate pattern classifiers

We used time-domain features in a multi-class classification pipeline with the EEG pattern from each of the four finger presses representing a class. We extracted time-domain features for classification, signals were smoothed using a moving averaging window of 100ms, wherein each time point is replaced by the mean of the 100ms around that point. This process was done for both sleep and wake data for each participant. Afterwards, sleep data (channels x time) from each participant is used in performing principal component analysis (PCA) and calculating the principal components. Following this, we calculated the explained variance for each principal component (eigen value of a component / sum of all eigen values), we then sorted the principal components (PCs) based on the explained variance and used the ones that contained 95% of the explained variance. Those PCs should be representing the dimensions in which the highest variance in the data exists and putative useful information. We then used the PCs and transformed both sleep and wake data which gave two transformed data sets containing PCs x time. We then used each time point observation to train our linear discriminant analysis (LDA) model (Blankertz et al., 2011). The trained LDA model was then applied to each time point after the cue in wake which yielded a classification accuracy at each wake time point. A classification output was then obtained from each participant and the final output was compared to chance level of 0.25. The result was then corrected for multiple comparisons using cluster-based permutation in Fieldtrip (Oostenveld et al., 2011) and lively vectors (lv)(Abdellahi, 2022) which gave the same results. For cluster-based permutation, Monte Carlo was used with a sample-specific test statistic threshold = 0.05, permutation test threshold for clusters = 0.05, and 100,000 permutations. The correction window was the whole length of wake trial (1.15 second).

### Compression and dilation of reactivation

A popular method for detecting the temporal compression of replay and used in the rodent literature is the template matching method. Generally, in template matching, a template is used from sleep episodes and this template is then slid on wake activity during maze navigation and a correlation coefficient is calculated which indicates the similarity of firing activity between the template and the window. This process is repeated for different scaling factors such that the windows are resized to smaller or longer sizes the process was repeated to measure compression and dilation of replay. The spatial resolution of EEG signals is low, however, signals measured at different channels in sleep can be compared to the same channels in wake to infer their degree of similarity at different compression/dilation ratios. In our data, we adopted a classification-based method to detect compression/dilation of reactivation given the differences between EEG of multiple classes with TMR and continuous firing pattern and that our classifiers can adapt to sleep and detect their subtle features. We thus used different temporal ratios that represent the ratio between sleep trial and wake trial and for each ratio we evaluated the classification performance. We tested different temporal ratios that ranged from 20 faster to 2.2 slower reactivation compared to wake. Theoretically, we could check for faster compressions given that classification was significant for the 20 times faster reactivation. However, we did not go beyond 20 times because the sliding window in sleep will be shorter than 10samples (50ms) and such very short window will not be reliable to resize and relate to wake to classify reactivation. In the meantime, we stopped at 2.2 slower reactivation because this matches the length of minimum sleep trial of 2.5seconds divided by the length of wake trial of 1.15seconds, thus, we stopped at this number to prevent any missing data points. We adopted a similar approach of calculating the PCA and transform the channels into PCs but we did not smooth the signals to keep the temporal information intact as smoothing could impact short effects. Both sleep and wake were transformed with the same PCs using sleep data, so the features are projected to the same feature space. This implies that if there was an activity on specific PCs in sleep the model will look at the same PCs in wake which will guarantee spatial alignment. Afterwards, for a specific compression ratio all sliding windows are used and resized to match the length of wake trial then all their data points are aggregated and used as features. Those aggregated features were then reduced with PCA using sleep data and the PCs containing 95% of explained variance were used to transform both sleep and wake data. Then LDA model was trained with sleep and tested with wake to yield a classification output for a particular compression ratio. This process was repeated for all compression ratios and all participants. Classification performance was then compared to chance level of 0.25 for each compression ratio using a Wilcoxon signed-rank test.

### Spindle analysis and spindle-based reactivation predictors

We analysed post-cue spindle activity to check if it relates to detected reactivation. We band-pass filtered our sleep data in the range [11 16]Hz using channel Cz and used the time duration [0 2.5]seconds then we used Hilbert transform. Afterwards, we used the instantaneous magnitude and phase that resulted from the Hilbert transformation to get the power by taking the absolute value of the complex vector to get the magnitude and then squaring that magnitude to get the power. We then divided our trials into two groups one with higher than median post-cue sigma power and the other with lower than median.

Following our prior work on the relationship between spindles and reactivation(Abdellahi et al., 2023a). We found that pre-cue spindle features can be used to predict TMR success in triggering reactivation and thus we had predictors that can differentiate correct/incorrect trials. Thus, we suggested that this can be used to guide the delivery of TMR cues to maximise reactivation. Here, we used the trained decision trees using data from all participants from our previous study and extracted the same features from the current dataset (a Boolean value with whether there is a spindle before the trial:1 or not:0). Afterwards, we applied the classification model on the current data. We then used the predicted class of the classifier: correct/incorrect to apply our current classification pipeline twice, once for each group. Only the group of trials that were predicted to be correct showed significant classification performance. This shows that the pre-cue spindle features can be used to predict successful reactivation and when to apply TMR to maximise reactivation.

EEG cleaning and other analyses (classification, compression/dilation, spindle analyses) were conducted with lively vectors (lv) (Abdellahi, 2022) toolbox developed by Mahmoud E. A. Abdellahi and it uses some functions from Fieldtrip (Oostenveld et al., 2011), MVPA-light (Treder, 2020), EEGLAB (Delorme & Makeig, 2004), and built-in Matlab functions.

## Data availability

Data and scripts are available on OSF: (https://osf.io/byvcg/?view_only=9b149e0387814bf1a6fca692f90e9167) including the EEG data, behavioural files, all the analyses with MATLAB scripts. Participants private identifications are all anonymized.

## Acknowledgements

This work was supported by the ERC grant SolutionSleep to P.A.L and ERC funded the Ph.D. of M.E.A.A. We would like to thank Niall McGinley and Ibad Kashif for helping in recruiting participants and scoring sleep data. We would like to thank Anne C. M. Koopman and all members of our group NaPs for their useful input.

## Author contributions

M.E.A.A., M. R. and P.A.L. conceptualisation and investigation of the experiment. M.E.A.A. and M. R. collected the data. M.E.A.A. analysed the data and wrote the original draft. M.E.A.A., M. R., P.A.L., and M.S.T reviewed and edited the manuscript.

