## Supplementary Material for "Targeted memory reactivation elicits temporally compressed reactivation linked to spindles"

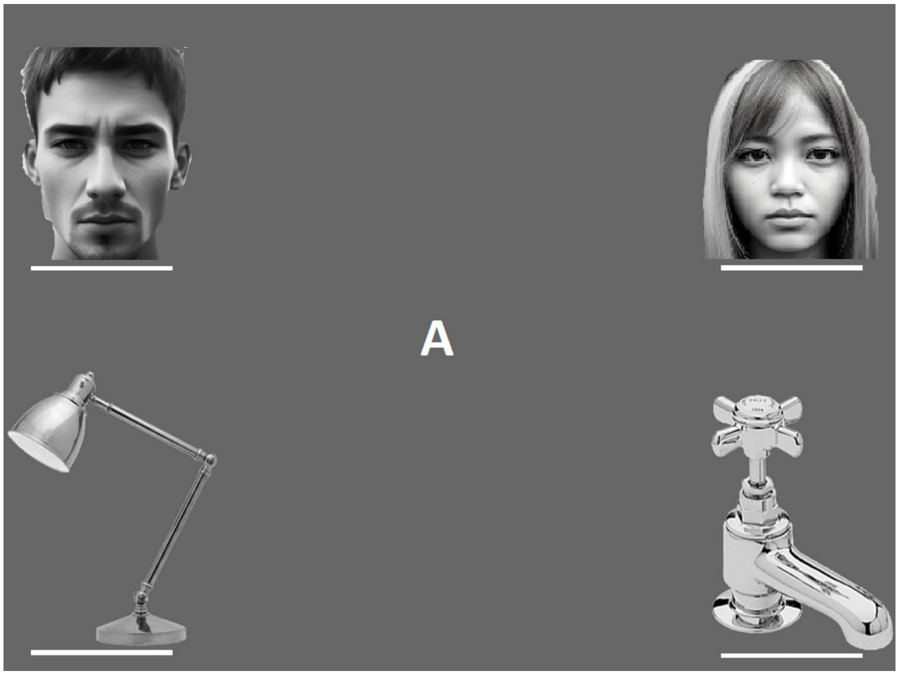


Supplementary figure 1: Illustration of the four images that appeared in the task: two faces and two objects. Faces in this illustration were artificially generated using AI.

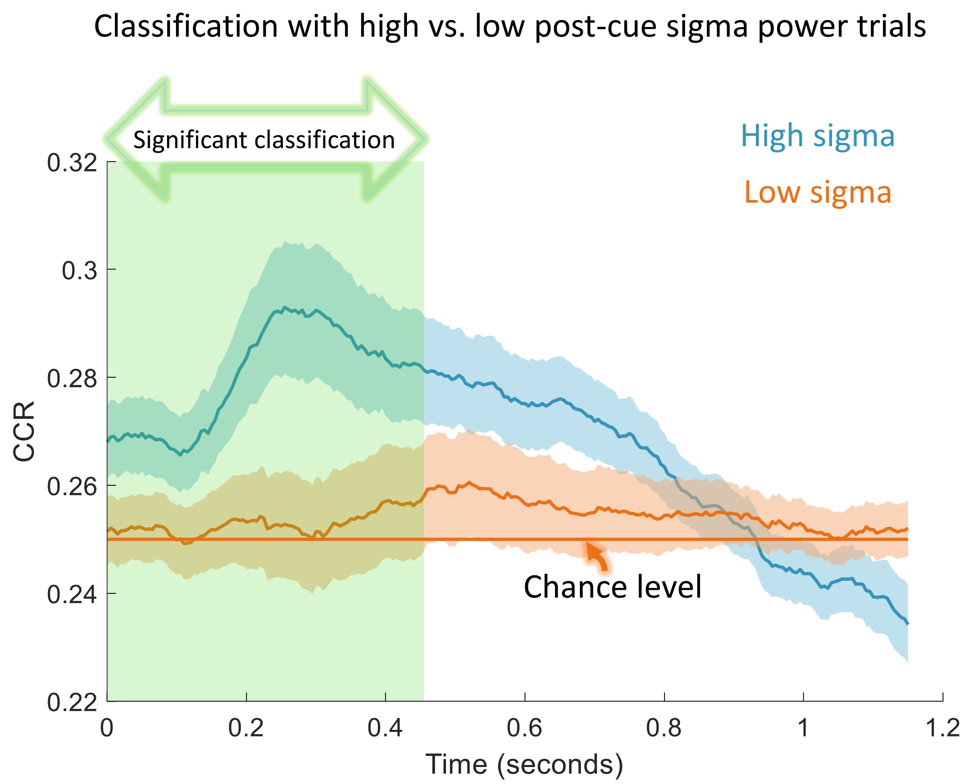


Supplementary figure 2: Classification performance for trials with high post-cue sigma power compared to trials with low post-cue sigma power. This shows a significant difference explained by the cluster shaded in green (n=48, p = 0.022).
